# Brain-blood biomarkers take a walk on the wild side: glial responses to environmental conditions and individual traits in wild frogs

**DOI:** 10.1101/2025.07.07.663561

**Authors:** Léa Lorrain-Soligon, Christophe J. Dubois, Coraline Bichet, François Brischoux, Jérôme Badaut

**Affiliations:** Centre d’Etudes Biologiques de Chizé, CEBC UMR 7372 CNRS – La Rochelle Université, 79360 Villiers en Bois, France; Sorbonne Université, Université PSL, EPHE, CNRS, Milieux Environnementaux, Transferts et Interactions dans les hydrosystèmes et les Sols, METIS, Paris, France

**Keywords:** Brain plasticity, Captivity, GFAP, Immunity, Osmolality, Sex-specific

## Abstract

Proteins from brain cells, including Glial fibrillary acidic protein (GFAP), has been tested for diagnostic and prognostic of neurological dysfunctions. Release of GFAP into the blood-stream, may be a consequence of its up-regulation in reactive astrocytes. However, astrocytic-GFAP expression is also increased during brain remodeling after physiological perturbations such as osmotic challenge. The presence and quantification of GFAP in blood circulation have never been investigated in the context of brain responses to environmental variations in wildlife. In a wild amphibian (green frogs*, Pelophylax* sp.), captured in several ponds with different salinity, GFAP was detected in plasma. Males from more saline ponds exhibited higher plasmatic GFAP levels, independent to their blood osmolality, suggesting that plasmatic GFAP-level reflects cerebral response to osmotic challenge. Plasmatic-GFAP correlated with immune markers (hemoglobin binding proteins, lymphocytes, neutrophils and monocytes), size and body condition, reinforcing its role as a physiological biomarker. We also highlighted that captivity had a significant effect on plasmatic-GFAP levels with sex-specific dynamics, masking the response to a short-term experimental salinity exposure. For the very first time, we show that plasmatic-GFAP levels could be a biomarker of brain plasticity to environmental conditions, physiological traits, and stress responses in wildlife.

**Significant Statement:** We investigated the use of brain Glial fibrillary acidic protein (GFAP), a cytoskeletal protein of astrocytes, that has been associated with brain disorders in clinical studies, as a biomarker of environmental conditions and individual traits in wildlife. In a wild amphibian, plasmatic GFAP-levels were correlated with the salinity of ponds, immune markers and size for males. GFAP-levels were correlated with body condition for both sexes. Interestingly, captivity induced transient increase of plasmatic-GFAP levels, probably due to stress. In our study, we demonstrate that plasmatic-GFAP levels may represent an excellent brain biomarker of its plasticity to environmental conditions, physiological traits, and stress responses in wildlife.

## 1. INTRODUCTION

During the past decade, tremendous number of studies have reported the discovery of new blood biomarkers for diagnosis and prognostication in various neurological indications, to overcome the cost and difficulties to perform neuroimaging and cerebrospinal fluid (CSF) biomarkers (1). These studies have focused on various proteins expressed by brain cells, including glial fibrillary acidic protein (GFAP), an intermediate filament protein expressed in cytoskeleton of the astrocytes (2). GFAP have been found to increase in various brain disorders, as a part of neuroinflammatory processes. Increased GFAP level in the plasma has been explained by the blood-brain interface alterations and release of GFAP by reactive astrocytes (for review (1)). However, human studies have reported baseline release of astrocytic proteins, including GFAP in physiological situation (3). To date, the mechanisms responsible for GFAP release in blood stream, and its functional consequences are still unknown.

Independent of neuroinflammatory processes, GFAP expression and distribution, and astrocyte morphological changes have also been described under physiological conditions, such as hydric stress (4–6). Neurons of the hypothalamus, including supraoptic and paraventricular nuclei, play a crucial role for hydric homeostasis and osmotic regulation in vertebrates (7–9). Indeed, magnocellular oxytocin (OT) and vasopressin (VP) positive neurons release neurohormones involved in osmotic response (4), but also in stress response (10). Interestingly, the magnocellular neuroendocrine system exhibits an exceptional structural plasticity, very well described in rodents, involving a full remodeling of the astrocytes (4, 11–13); including significant morphological changes facilitating the neuronal firing synchronization for release of OT and VP in systemic blood circulation in pituitary gland (Fig 1A) (4, 5). This astrocyte remodeling is correlated with the alteration of GFAP expression in the hypothalamic nuclei (6).

**Figure 1.**
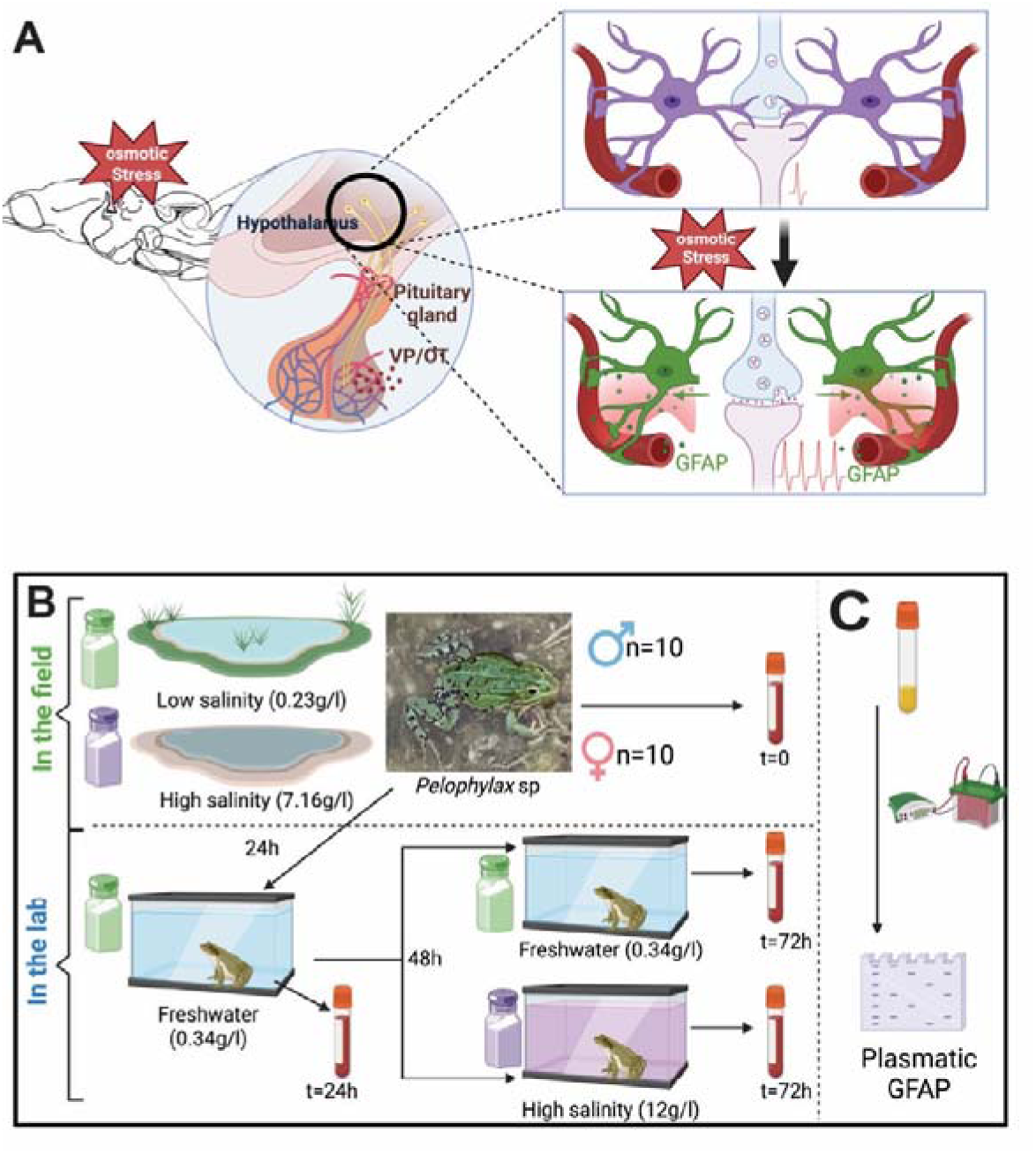
(A) Schematic representation of astrocyte remodeling and GFAP decrease after an osmotic stress in the brain hypothalamus. Osmotic stress induces a decrease of astrocytic coverage at the synaptic clefts, facilitating the synchronization of the neuronal activation for the release of neurohormones, oxytocin (OT) and vasopressin (VP), in the general blood circulation by the pituitary gland (B) schematic representation of the experimental protocol regarding exposure of adult *Pelophylax* sp. (green frogs) in the field and in the lab and (C) of computation of plasmatic GFAP levels.

Environmental conditions shape animals physiological responses, influencing their ability to adapt and survive in changing environments (14, 15). Despite a large literature on physiological responses to environmental variations (16), very little is known about the consequences of these stressors on brain functions in wildlife, due to limited noninvasive readouts. Among these stressors, increasing salinity, linked to anthropogenic disturbances such groundwater pumping (17), use of de-icing salt (18), reduction in freshwater flow and droughts (17), and sea level rise (19), present significant challenges, particularly for aquatic organisms. Indeed, most of them need to maintain homeostasis (20–22). Amphibians frequently experience variable salinity, and osmoregulatory strategies are critical for their survival (23, 24). Saline exposure induce a set of physiological responses, often modulated by sex (25), as males and females differ in their investment in reproduction and metabolic demands (26, 27), leading to sex-specific reproductive physiological trade-offs. Plasma osmolality is a common physiological marker for osmotic stress recorded for amphibians (25, 28, 29), however it provides only a snapshot of an individual’s current physiological state. Additionally, the physiological mechanisms of osmoregulation are well-documented (22), but less known is how environmental salinity influences brain plasticity and stress responses. To the best of our knowledge, changes in the plasmatic level of GFAP have never been investigated in such physiological conditions in vertebrates.

Using a coastal amphibian (green frogs, *Pelophylax* sp.) naturally exposed to variable salinity as a study model, we combined field and experimental approaches to comprehensively test the hypothesis that GFAP can be released in the blood circulation during brain astrocyte remodeling in response to osmotic stress (Fig. 1). We predicted that plasmatic GFAP levels can be used to test the effects of osmotic stress (salinity) on brain responses and can correlate with immune markers identified as a response to increased osmotic stress in amphibians (25, 30).

## 2. METHODS

### 2.1. Field procedure and housing conditions

The study was carried out on the “Réserve Naturelle Nationale de Moëze-Oléron” (45°53’33.36"N, 1°04’59.16"W), located in the Atlantic coast of France (Charente-Maritime) where Green frogs (*Pelophylax* sp.) are widespread (31, 32). In this study area, *Pelophylax* sp., are composed of viable and fertile hybrids (Graf’s hybrid frog, *P. kl. grafi*) of the Marsh frog (*P. ridibundus*) and the Perez’s frog (*P. perezi*, (33)). This species is primarily aquatic and is relatively tolerant to salt (34–36). In our study site, *Pelophylax* sp. can be found in ponds in which salinity ranges from 0.14 g.l^-1^ to 16.19g.l^-1^ (31).

Male (n=10) and female (n=10) adults were captured at night using a net, between 10pm and 12pm from 9 different ponds (each separated by less than 1.3 km from one another, salinity ranging from 0.23 g.l^-1^ to 7.16 g.l^-1^ [mean= 2.887 ± 0.488 g.l^-1^]) from 30 March 2021 to 29 May 2021 (Fig.1B). Males were found in ponds characterized by a wider range of salinity than females (respectively 0.61-7.16 g.l^-1^ [mean 3.854 g.l^-1^], 0.23-4.71 g.l^-1^ [mean 1.919 g.l^-1^]). After capture, individuals were brought to the laboratory, where they were weighted (with an electronic balance ±0.1 g, Ohaus) and measured for body size (snout-vent length: SVL) using a calliper (±0.1mm). All captured individuals weighed more than 15g, allowing an accurate determination of the sex, based on the presence of secondary sexual characteristics.

After these measurements (within 2 hours after capture), we collected blood samples (∼1% body mass) through cardiocentesis, using 1 ml syringes and heparinized 30G needles. A drop of blood was smeared on a slide for further leukocytes count (see 2.3.3). Remaining collected blood was centrifuged (7 min at 2000 G) to separate red cells from plasma. Plasma fractions were collected and stored at −20 °C until further analyses of plasma osmolality, hemoglobin-binding proteins, and GFAP levels (see 2.3.1 and 2.3.2).

After cardiocentesis, frogs were placed individually in transparent plastic boxes (14 × 16 × 9 cm) filled with water (see details below) with a perforated cover. They were kept in a room with natural photoperiod, and were not fed during the duration of the experiment. At the end of the experiment, they were released to the pond from which they were captured.

### 2.2. Experimental design

Two successive stages were performed in our experiment (see (25) for details). First, boxes were filled with freshwater (0.34 g.l^-1^ salinity ± 0.007 SE) for 24 h to allow frogs to restore osmotic balance (hereafter termed “acclimation”). Individuals were then randomly allocated to the experimental groups. In this second stage, (hereafter termed “Treatment”), frogs were subjected for 48 hours to either freshwater (0.35 g.l^-1^ salinity ± 0.003 SE, n=5 males and n=5 females) or 12 g.l^-1^ (12.08 g.l^-1^ salinity ± 0.040 SE, n=5 males and n=5 females). Water was prepared by dissolving sea salt in tap water. A blood sample was collected and a blood smear performed at the end of the acclimation period (24 hours post-capture) and at the end to the treatment period (72 hours post-capture), as described above, to compute for changes in plasma osmolality, leukocytes count and GFAP levels.

### 2.3. Blood samples analyses

#### 2.3.1. Western blots and Analysis for plasmatic GFAP

After protein assay (BCA kit, Thermofisher, 23227), samples (30µg/lane) were loaded on SDS-PAGE gels (Thermofisher NW04120BOX) following vendor’s protocol. Then, proteins were transferred to nitrocellulose membranes (Thermofisher IB24002). Membranes were blocked in blocking buffer (LiCor, Lincoln, NE, 927-75001) prior to overnight incubation with primary antibody at 4°C. All antibody incubations were in blocking buffer with 1% tween-20. The primary antibody used was chicken anti-GFAP (1:3,000, Abcam, AB4674). Membranes were washed in PBS with 0.1% tween-20 and then incubated at room temperature for 1 hr with a donkey-anti-chicken secondary antibody coupled with IRDye680 (1:20, 000, LICOR, 926-68028). Membranes were finally washed, and bands were visualized using the LiCor infrared Odyssey imager (LiCor, Lincoln, NE, Fig.2A). Protein density quantification was performed using Image J software (NIH) and target protein levels were normalized to the corresponding loading control levels.

**Figure 2.**
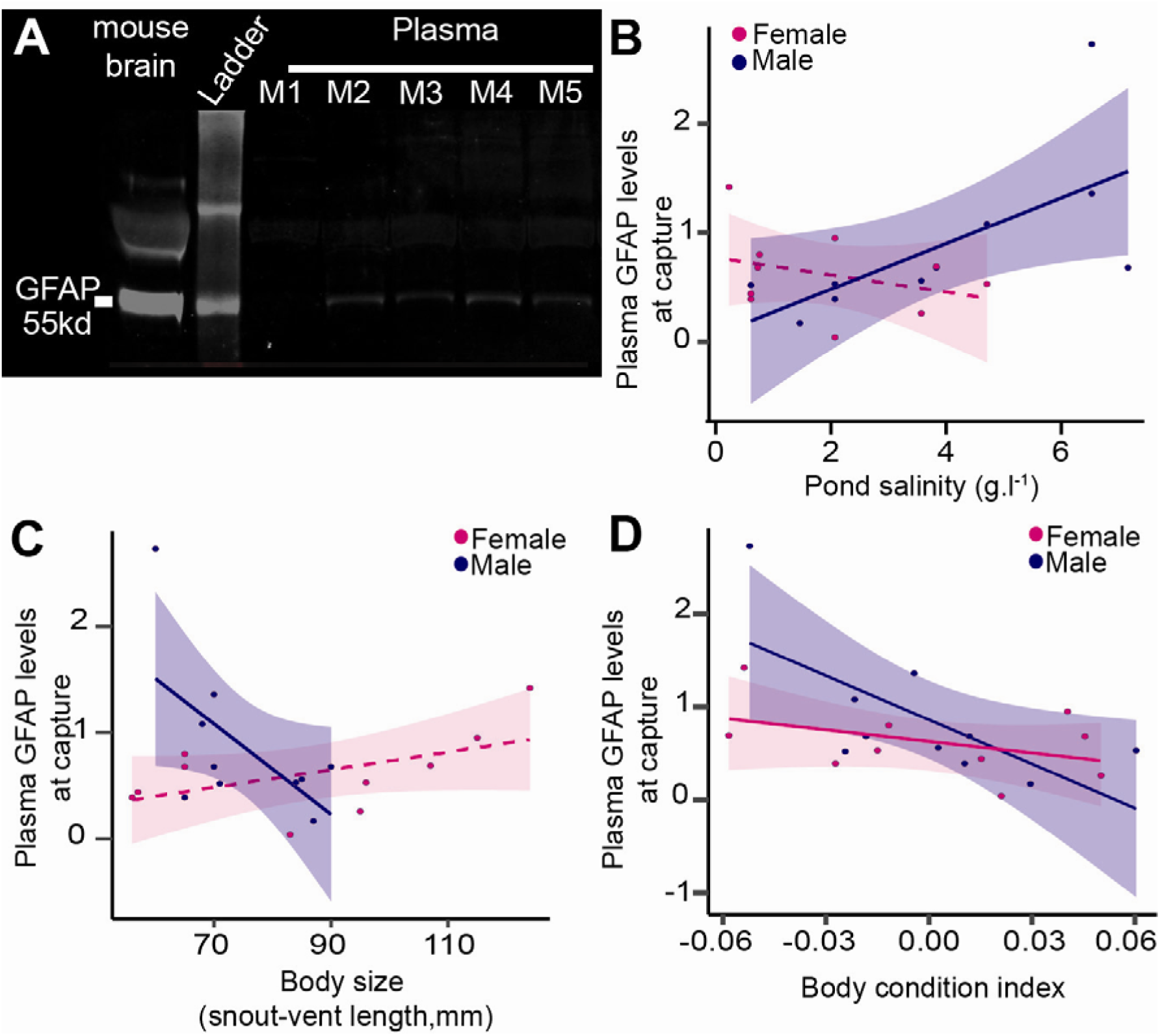
(A) Western blot example of GFAP in male green frog plasma (M1, M2, M3, M4, M5) from different ponds showed the presence of GFAP in plasma with 55kDa, similar to size of GFAP detected in mouse brain. Intensity of the band demonstrated various plasmatic levels for GFAP protein. (B) Effect of pond salinity, (C) Snout-Vent Length (SVL, body size), and (D) body condition index on plasmatic GFAP levels at capture. Solid lines represent a significant relationship at α=0.10, dashed lines represent no significant relationship.

#### 2.3.2. Osmolality and hemoglobin-binding proteins

Plasma osmolality (mOsmol.kg^−1^) was measured from 10 μl aliquots on a Vapro2 osmometer (Elitech group). We quantified the hemoglobin-binding proteins (mg.ml-^1^) of frogs in 7.5 µl plasma samples using a commercially-available colorimetric assay (TP801, Tri-Delta Diagnotics, USA). We followed the instructions provided by the manufacturer with slight modifications following Matson et al. (2012) (see (25)).

#### 2.3.3. Leukocyte profiles

Blood smears were air-dried and fixed in absolute methanol for 5 min, and then stained using May-Grünwald-Giemsa staining method (3 min in May-Grunwald working solution; 1 min in pH 7 buffer, and 10 min in Giemsa solution diluted at 1/20 in the pH 7 buffer). Neutrophils, lymphocytes, monocytes and eosinophils were counted by a single observer (LLS) up to 100 leukocytes (the number of each leukocyte type for 100 leukocytes was then used), using a X100 ocular microscope (Microscope Primo Star 8, total magnification X1,000) and immersion oil, to determinate leukocyte profiles.

### 2.4. Statistical analyses

All statistical analyses were performed using R 3.6.3 (38) and Rstudio v1.1.419. We computed Linear Models (LMs), Gamma Generalized Linear Models (GLMs) (Table 1) or Linear Mixed Models (LMMs), Gamma Generalized Linear Mixed Models (GLMMs) (Table 2), using the lme4 package (39). For LMs and LMMs, normality was checked (shapiro test) and data were log10 or sqrt transformed when residuals did not follow the normality assumption. For all computed tests, models accuracy was also tested using the check_model function from the *performance* package (40). Significance level was set at α<0.05, effects were considered trending significant at α<0.10. Best variables were retained using a top-down selection procedure.

**Table 1.** Statistical results for the effect of environmental and physiological variables (Pond salinity, osmolality, SVL, body condition index, hemoglobin-binding protein, neutrophils, lymphocytes, monocytes and eosinophils, as well as their interaction with sex) on GFAP levels at capture, including the type of model used. Best models were obtained using a top-down selection process.

| Factors | Model | Sum Sq | F value | p-value |
| --- | --- | --- | --- | --- |
| Pond salinity |  | 0,972 | 3,955 | **0,064** |
| Sex | LM | 0,003 | 0,010 | 0,921 |
| Pond salinity*Sex |  | 1,314 | 5,344 | **0,034** |
| Osmolality |  | <0,001 | 0,001 | 0,983 |
| Sex | LM (log10 transformed) | 0,127 | 0,784 | 0,389 |
| Osmolality*Sex |  | <0,001 | 0,001 | 0,977 |
| SVL |  | 0,003 | 0,010 | 0,922 |
| Sex | LM | 0,301 | 1,189 | 0,291 |
| SVL*Sex |  | 2,172 | 8,580 | **0,010** |
| Body condition index | LM (log10 transformed) | 0,034 | 6,855 | **0,017** |
| Hemoglobin-binding protein |  | 0,964 | 2,688 | 0,121 |
| Sex | Gamma GLM | 0,566 | 1,577 | 0,228 |
| Hemoglobin-binding protein*Sex |  | 2,659 | 7,412 | **0,015** |
| Neutrophils |  | 0,152 | 1,379 | 0,257 |
| Sex | LM (log10 transformed) | 0,099 | 0,898 | 0,358 |
| Neutrophils*Sex |  | 0,683 | 6,185 | **0,024** |
| Lymphocytes |  | 2,130 | 10,548 | **0,005** |
| Sex | LM | 0,270 | 1,337 | 0,265 |
| Lymphocytes*Sex |  | 0,860 | 4,258 | **0,056** |
| Monocytes | LM (sqrt transformed) | 0,410 | 5,282 | **0,034** |
| Eosinophils |  | 0,037 | 0,353 | 0,561 |
| Sex | LM (sqrt transformed) | 0,052 | 0,500 | 0,490 |
| Eosinophils*Sex |  | 0,021 | 0,202 | 0,659 |

**Table 2.** Statistical results for the variations of GFAP according to the sex of individuals and period of sampling, either after acclimation or after treatment. Best models were obtained using a top-down selection process.

| Period | Factors | Model | Chisq | p-value |
| --- | --- | --- | --- | --- |
| After<br>acclimation<br>(24 hours<br>captivity) | Sex | Gamma<br>GLMM | 0.422 | 0.516 |
|  | Period |  | 1.903 | 0.168 |
|  | Sex*Period |  | 6.387 | **0.012** |
| After<br>treatment<br>(72 hours<br>captivity) | Sex | LMM | 0.101 | 0.751 |
|  | Period |  | 1.103 | 0.319 |
|  | Sex*Period |  | 15.757 | **<0.001** |
|  | Treatment |  | 0.121 | 0.728 |
|  | Sex*Treatment |  | 0.058 | 0.810 |

**Table 3.**
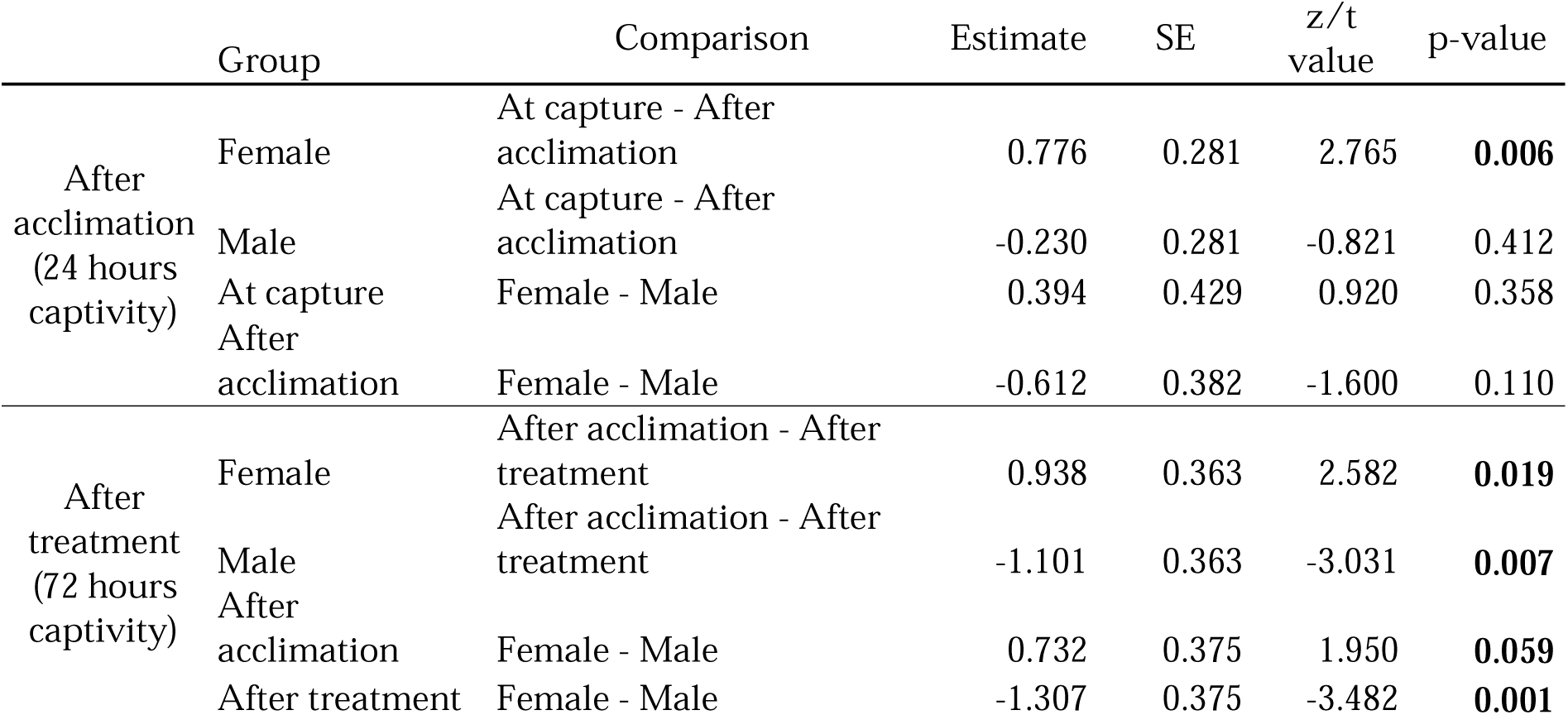
Post-hoc results for the variations of GFAP according to the sex of individuals and period of sampling, either after acclimation or after treatment. Best models were obtained using a top-down selection process.

#### 2.4.1. Variability at capture

From the samples and measures took at capture, we tested for the variation of GFAP according to pond salinity, osmolality, SVL, body condition index [as the residuals of the relation between log10(mass) and log10(size)], hemoglobin-binding protein, neutrophils count, lymphocytes count, monocytes and eosinophils count (as well as their interaction with sex) (see table 1 for models used). Given the low count of individuals and to avoid autocorrelation, we ran separate models for each explanatory variable, including the interaction with sex. The correlation between pond salinity and osmolality, a common marker of exposure to salt, was also tested using a LM in order to determine if an increased salinity reflect a higher osmolality in our sample.

#### 2.4.2. Experimental variability

After acclimation, we tested for the variation of GFAP according to period of sampling (at capture or after acclimation), sex, and their interaction using a Gamma GLMM. After treatment, we tested for the variation of GFAP according to the period of sampling (at capture or after treatment), sex, treatment (0 or 12 g.L^-1^ salinity treatment) and their triple interaction using a LMM. We also computed Pearson correlation tests between GFAP levels at capture and after acclimation, at capture and after treatment, and after acclimation compared to after treatment. Finally, using the same model structure as in 2.4.1, we tested for the variation of GFAP according to pond salinity, osmolality, hemoglobin-binding protein, neutrophils count, lymphocytes count, monocytes and eosinophils count as well as their interaction with sex) after acclimation and after treatment (see Appendix A for models used).

## 3. RESULTS

### 3.1. Plasmatic-GFAP characterization and quantification in wild amphibians

GFAP was detected in all the plasma sampled with expected molecular weight at 55kda, as observed for mouse (Fig2A). Various intensities were observed in males from different ponds with different salinities (Fig2A). From the blot, intensity of the GFAP band was quantified for each individual.

### 3.2. GFAP response to environment (pond salinity) at capture

At capture, plasmatic-GFAP levels varied with the interaction between pond salinity and sex (Figure 2B, Table 1). Specifically, plasmatic GFAP levels increased with pond salinity, but only in males (Estimate=0.210 ± 0.084, t-value=2.504, p-value=0.037) while it was not related to pond salinity in females (Estimate=-0.080 ± 0.080, t-value=-0.986, p-value=0.353). However, GFAP levels did not correlate with osmolality, nor its interaction with sex (Table 1). Osmolality and pond salinity were marginally correlated (Estimate=4.270 ± 2.310, t-value=1.849, p-value=0.081).

### 3.3. GFAP response to morphology at capture

Plasmatic GFAP levels varied with the interaction between SVL (body size) and sex (Figure 2C, Table 1). Specifically, GFAP levels marginally decreased with SVL, but only in males (Estimate=-0.043 ± 0.020, t-value=-2.155, p-value=0.063) while it was not related to SVL in females (Estimate=0.008 ± 0.005, t-value=-1.784, p-value=0.112). Plasmatic GFAP levels decreased for both sexes with increasing body condition (Figure 2D, Table 1).

### 3.4. GFAP association with immune parameters at capture

Plasmatic GFAP levels varied with the interaction between hemoglobin-binding proteins concentration and sex (Figure 3A, Table 1). GFAP levels increased with hemoglobin-binding proteins concentration in males, (Estimate=5.200 ± 1.145, t-value=4.542, p-value=0.002), but not in females (Estimate=-1.223 ± 0.842, t-value=-1.453, p-value=0.184).

**Figure 3.**
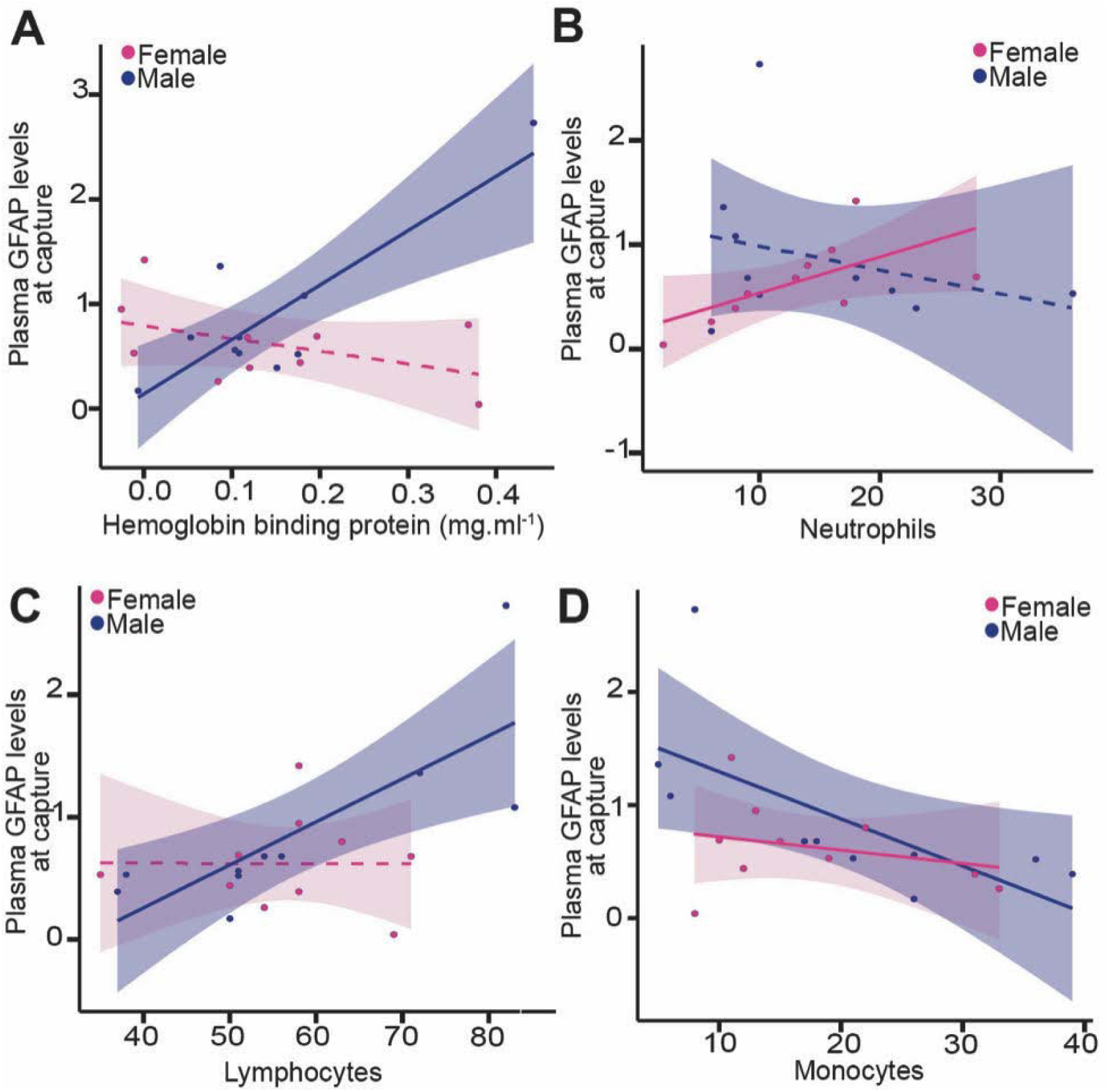
Effect of (A) hemoglobin-binding protein, (B) neutrophils, (C) lymphocytes and (D) monocytes on plasmatic GFAP levels at capture. Solid lines represent significance at α=0.10, dashed lines represent no significant relationship.

GFAP was also found to be related with leukocyte profiles. GFAP levels varied both with the interaction between neutrophils number and sex and with the interaction between lymphocytes number and sex (Figure 3B, C, Table 1). GFAP levels increased with the number of neutrophils in females, (Estimate=0.034 ± 0.015, t-value=2.239, p-value=0.056), but not in males (Estimate=-0.023 ± 0.026, t-value=-0.881, p-value=0.404). Conversely, GFAP levels increased with the number of lymphocytes in males (Estimate=0.035 ± 0.010, t-value=3.565, p-value=0.007), but not in females (Estimate=-0.002 ± 0.013, t-value=-0.019, p-value=0.985). Moreover, GFAP levels positively correlated with the number of monocytes in both sex (Figure 3D, Table 1). GFAP was not related to the number of eosinophiles (Table 1).

### 3.5. Effects of captivity on plasmatic GFAP levels

Acute salinity treatment (12 g.l^-1^) did not have any effects on plasmatic GFAP levels for both sexes (Fig. 4A, Table 2). Interestingly, GFAP levels were significantly higher after acclimation (24h post-capture) than upon capture, but only in females (Figure 4B, Tables 2,3), while it remains constant after acclimation in males. GFAP levels were higher in males after treatment (72h post-capture) than after acclimation (24h post-capture), while in females GFAP levels were lower after treatment than after acclimation, independently of experimental salinity exposure for both sexes (Figure 4B, Tables 2,3).

**Figure 4.**
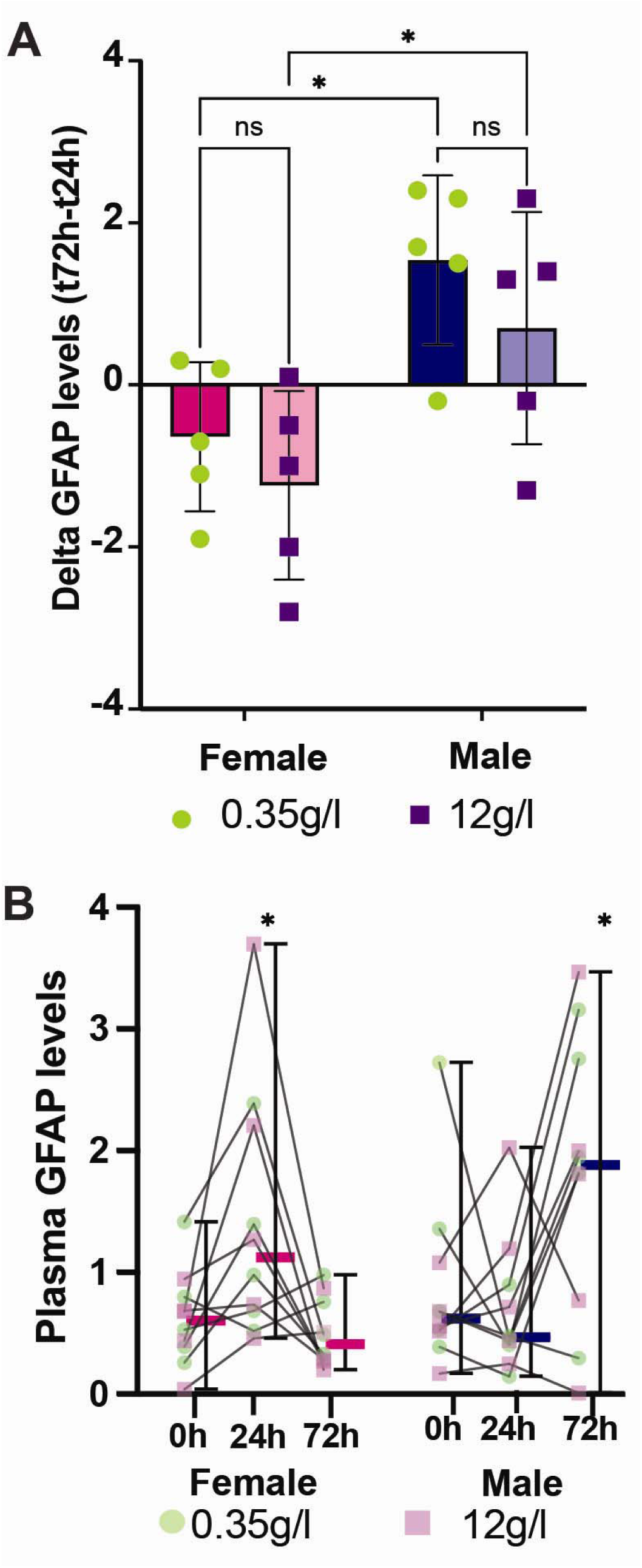
(A) Differences in plasmatic GFAP levels (Mean **±** SE) across the experimental treatment (exposure at 0 or 12 g.L^-1^ salinity between 24h and 72h post-capture) according to the treatment and separately for each sex. (B) Plasmatic GFAP levels (Mean **±** SE) at capture, after acclimation (24h post-capture) and after treatment (0 or 12 g.L^-1^ salinity, 72h post-cpature) according to the treatment and separately for each sex.

GFAP levels were higher in females compared to males after acclimation, but higher in males than in females after treatment (Figure 4B, Tables 2,3). GFAP levels between capture and acclimation (cor=0.050, t=0.210, df=18, p-value=0.836), between capture and treatment (cor=0.226, t=0.985, df=18, p-value=0.338), and between acclimation and treatment (cor=-0.220, t=-0.959, df=18, p-value=0.350) did not correlate.

Additionally, after acclimation (24h post-capture) and after treatment (72 hours post-capture), GFAP levels were no longer correlated with pond salinity or any neuroinflammatory or immunological parameters (Appendix A).

## 4. DISCUSSION

For the very first time, we successfully detected plasmatic GFAP and quantified its level in a wild amphibian species in relation to environmental conditions (pond salinity) and individual traits (body condition, hemoglobin-binding proteins levels and leukocyte counts). We demonstrate the potential use of GFAP as a brain-blood biomarker of physiological stress in a non-model organism. Additionally, GFAP levels were influenced by captivity more than by salinity exposure challenge in experimental conditions.

### 4.1. Detection of GFAP in plasma from amphibians

To the best of our knowledge, this is the first report of circulating plasmatic-GFAP in non-model organisms. Like mammalian model organisms, amphibian astrocytes express GFAP as detected by immunolabelling (41). While the presence of GFAP in amphibians genome is discussed (42), immunolabelling targeting GFAP remains consistent with - and specific to - astrocyte cytoskeleton (41, 42). Here, immunoblot exhibited one GFAP-band at the expected molecular weight (55kda, Fig 2A); suggesting the presence of cytoskeleton proteins from astrocytes in blood circulation of wild frogs under wild conditions. Our results are in line with previous studies showing the presence of brain proteins in blood circulation (GFAP, S100, a Ca2+ binding protein family) under physiological situations and independently of brain injuries in both rodents and humans (3, 43, 44). Even if most studies have focused on the quantification of these plasmatic brain proteins in clinical studies (45), previous works and the present study raise the question about the mechanisms involved in the release and diffusion of these proteins in the blood stream (Fig. 1A) in absence of brain disorders, and its functional implications.

### 4.2. Plasmatic GFAP biomarker of cerebral plasticity responses to environmental conditions and individual traits

Plasmatic GFAP levels at capture were influenced by the salinity of the pond in which individuals were captured, but only in males. Osmotic stress due to increased salinity has been shown to induce significant astrocytic remodeling (4, 5) and GFAP redistribution to facilitate the synaptic contact between magnocellular neurons for release of VP (Fig 1A, (4, 5)). GFAP is mostly expressed in astrocytes, and at low level in peripheral cells, such as Schwann cells (46). The positive relationship between plasmatic GFAP levels and environmental salinity strongly suggests that plasmatic GFAP levels can mirror brain response in amphibians. As such, plasmatic GFAP might be used as a marker of physiological tolerance to salinity in relation to brain remodeling as described from experimental studies (6). This is corroborated by the fact that plasmatic GFAP is overall more elevated in male green frogs as compared to females, which are often found in more saline ponds (25), presumably due to higher costs of reproduction in females (26, 27), leading females to avoid this osmotically (and metabolically) challenging environment (22). Males from saltier ponds may exhibit naturally higher GFAP levels due to long-term exposure to osmotic challenges. While some studies found differences in brain GFAP expression between sexes (47), we only found differences through interactive effects, suggesting that the relationships between plasmatic GFAP, environmental and physiological characteristics are modulated by sex-specific responses.

Morphological characteristics were also shown to be linked to plasmatic GFAP levels. Body size (SVL) was negatively associated with GFAP levels in males, and thus larger individuals had lower plasmatic GFAP levels. Because larger size is associated with older individuals in amphibians (48), this suggest that older males may display a greater physiological capacity to buffer environmental stressors, which dovetails with the fact that larger individuals are found in saltier ponds (25). Alternatively, this may suggest a selective process through which only the more stress-resilient individuals grow to larger sizes. Importantly, brain-GFAP expression is also known to vary with age in rodent and humans (49), usually associated with increase of GFAP-positive astrocytes with increasing age (50). Conversely, there was no correlation between GFAP and size in females, suggesting that females of all size and/or age (considering that we only sampled adults) avoid challenging habitats, probably due to their higher energetic investment in reproduction (26, 27). Additionally, body condition was negatively correlated with GFAP in both sexes, indicating that individuals in poorer condition may be less resistant to stress (51, 52). This reinforces the idea that plasmatic GFAP is a sensitive biomarker of physiological state, responding not only to environmental factors but also to individual traits and/or physiological mechanisms.

Indeed, the observed correlations between plasmatic GFAP and immune markers reinforce its potential as a brain-stress biomarker, especially in males. The positive correlation between plasmatic GFAP and hemoglobin-binding proteins in males suggests a link with inflammation (53). In fact, the role of hemoglobin-binding proteins is to bind free hemoglobin during inflammation caused by reactive oxygen components and/or by the immune response (54, 55). Interestingly, the increase in plasmatic GFAP with increasing lymphocyte counts in males, rather than neutrophils, is not the classical pattern expected in acute stress or primary infection. While neutrophils typically increase during immune responses (56, 57), as shown for example in amphibians exposed to salt (25), an increase in lymphocytes number may reflect a different process, such as immune memory development or specific immune cell remodeling (58). Still, in both sexes, we observed that plasmatic-GFAP levels decreased with an increasing number of monocytes. Monocytes serve as phagocytic and antigen-presenting cells (59), and are the main phagocytic cells in vertebrates (60). Importantly, there is now information that monocytes respond to different environments by modulating gene expression and cytokine production allowing homeostasis maintenance (61). Increased monocyte number might indicate homeostasis reestablishment, although this relation remains to be assessed. Overall, individuals with higher plasmatic GFAP levels exhibit distinct leukocyte profiles that appear to differ between sexes. While in females, the positive correlation with neutrophils may point toward a stress-related pattern, in males, the association with lymphocytes may indicate immune system remodeling rather than acute stress. The increase in monocytes in both sexes further supports the idea of systemic immune involvement.

### 4.3. Impact of captivity and no effect of salinity treatment on plasmatic GFAP

Unexpectedly, we did not find an influence of experimental exposure to salinity (0 or 12 g.L^-1^ salinity) on plasmatic GFAP levels, in contrast to the effects of environmental salinity on plasmatic GFAP levels. Additionally, we did not find, after acclimation and treatment, relationships between the measured physiological variables and plasmatic-GFAP levels. However, captivity seemed to affect plasmatic-GFAP, suggesting that capture and captivity have stronger impact on plasmatic-GFAP levels than short-term exposure to hyperosmotic conditions. In fact, capture (62, 63), translocation (64) and captivity (65, 66) are all known to be stressful for wild animals. Similarly to hydric stress, acute stress caused by capture and captivity may act on hypothalamic-pituitary adrenal axis (HPA) with astrocyte and GFAP remodeling. We thus suggest that the variations in plasmatic-GFAP levels observed are linked to stress associated with these different phases, including acclimation to captive conditions and may explain cerebral glial plasticity. Together with the fact that plasmatic GFAP levels upon capture, after acclimation and after treatment were not correlated, this reinforces the fact that plasmatic-GFAP level is a plastic response in amphibians (49, 67).

However, these responses varied with sex. Indeed, it is often found that stress linked to captivity is sex-dependent in a large number of species (68, 69), which our results seem to suggest for amphibians too. After 24 hours of captivity (acclimation), females showed an increase in GFAP levels, while GFAP of males remained stable. This suggests that females may experience a greater initial stress response to captivity, potentially due to differences in behavioral or metabolic adjustments.

However, after longer captivity (72 hours in captivity independently of the experimental treatment), the pattern reversed with an increase of plasmatic GFAP levels in males but a decrease of plasmatic GFAP levels in females. This pattern suggests that males may take longer to activate stress response, whereas females show a faster, but transient, response. The reversal of GFAP patterns over time further supports the idea that plasmatic-GFAP levels are highly plastic and responsive to environmental conditions, as well as a function of individual characteristics, highlighted by sex-specific responses.

## 5. CONCLUSION

Our study shows, for the first time, that plasmatic GFAP is detectable in the plasma of non-model wildlife species (here amphibians), and seems to be an excellent biomarker of brain plasticity to environmental conditions, physiological traits, and stress responses, with sex-specific dynamics. GFAP appears to be a highly plastic brain-blood biomarker, responding dynamically to environmental salinity, immune parameters, and captivity. Further studies should explore whether plasmatic GFAP levels are predictive of long-term survival and fitness in natural populations, and assess the generality of these patterns in a broader range of wild vertebrate species to better understand its ecological and physiological significance. Our observations also opened the biological mechanisms in GFAP release and physiological consequences in “brain-body” interactions.

### Ethic statement

This work was approved by the French authorities under permits R-45GRETA-F1-10, 135-2020 DBEC and APAFIS# 30169.

## Acknowledgments

The authors would like to thank all the staff of the Moëze-Oléron reserve (Philippe Delaporte, Pierre Rousseau, Vincent Lelong, Nathalie Bourret, Emma Bezot-Maillard, Loïc Jomat, Stéphane Guenneteau, Eliott Huguet and Julia Guerra Carande) for their welcome during field session. The authors thank Mathilde Malinconi for running western blot experiments, and Frédéric Angelier as co-PI of the NeuroEco Project (CNRS-MITI).

## Author contributions statement

LLS, FB and JB have conceptualized the study. LLS performed field and experimental settings, CD, CB, and JB participated to laboratory measurements. LSS, FB and JB participated to data analyses. LLS, FB and JB wrote the initial draft. FB and JB contributed equally to this manuscript and must be considered co-last authors. All authors have reviewed and edited the manuscript and approved the final version.

## Funding

Funding was provided by the CNRS, CNRS-MITI (NeuroEco), La Rochelle Université, the LPO, the Agence de l’Eau Adour-Garonne, the Conseil Départemental de la Charente-Maritime, the ANR (ANR-18-CE32-0006 [PAMPAS] and ANR-Eranet Neuvasc), the Beauval Nature association and the Région Nouvelle Aquitaine (Projet d’Observatoire du Marais de Brouage - PSGAR CRNA 2025). This study is part of the long-term Studies in Ecology and Evolution (SEE-Life) program of the CNRS.

### Data Availability Statement

Data will be made available upon acceptance.

## Appendix captions

Appendix A - Statistical results for the effect of environmental and physiological variables (Pond salinity, osmolality, hemoglobin-binding protein, neutrophils, lymphocytes, monocytes and eosinophils, as well as their interaction with sex) after acclimation (24 hours in captivity) and treatment (72 hours captivity), including the type of model used.

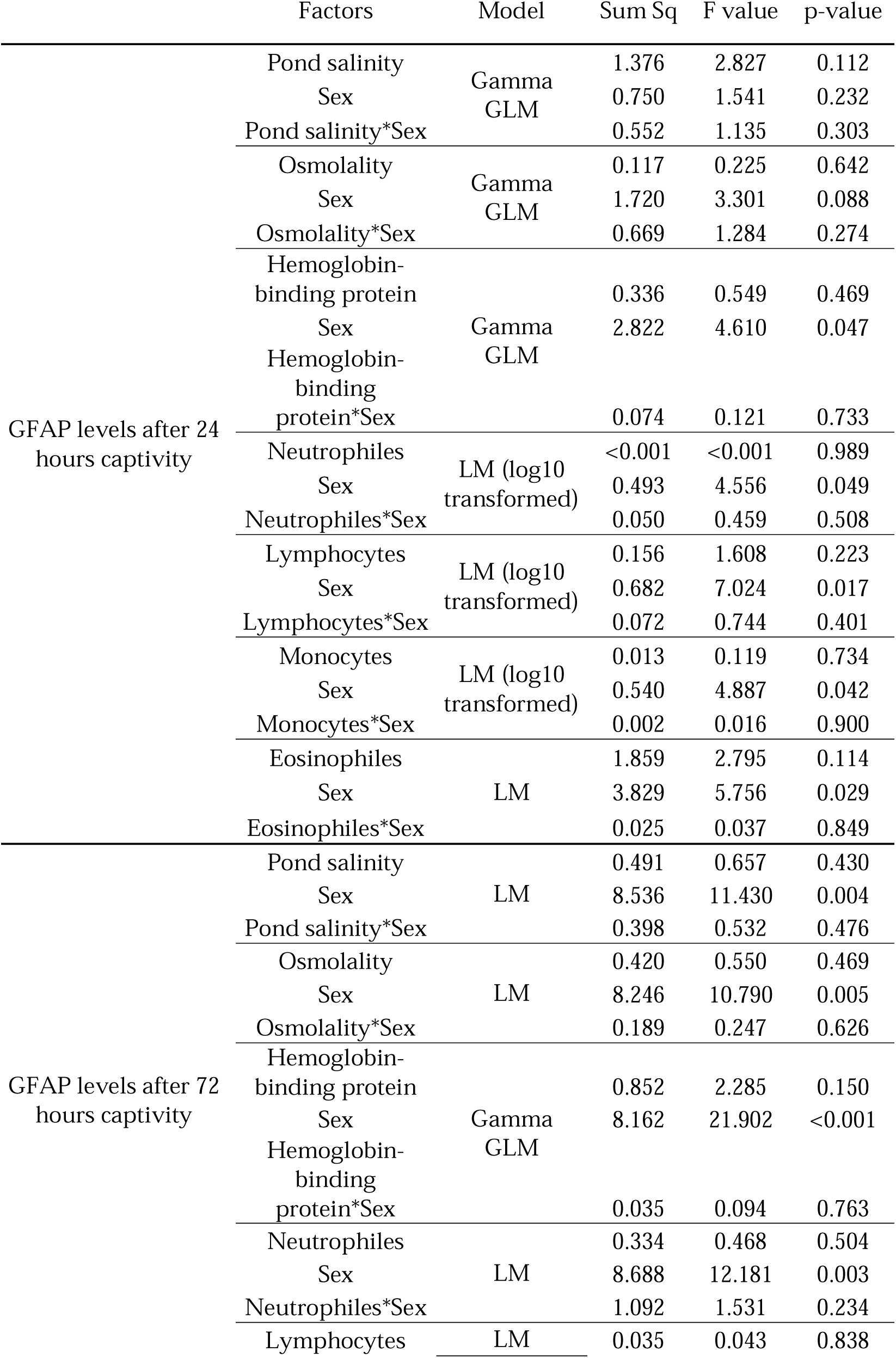
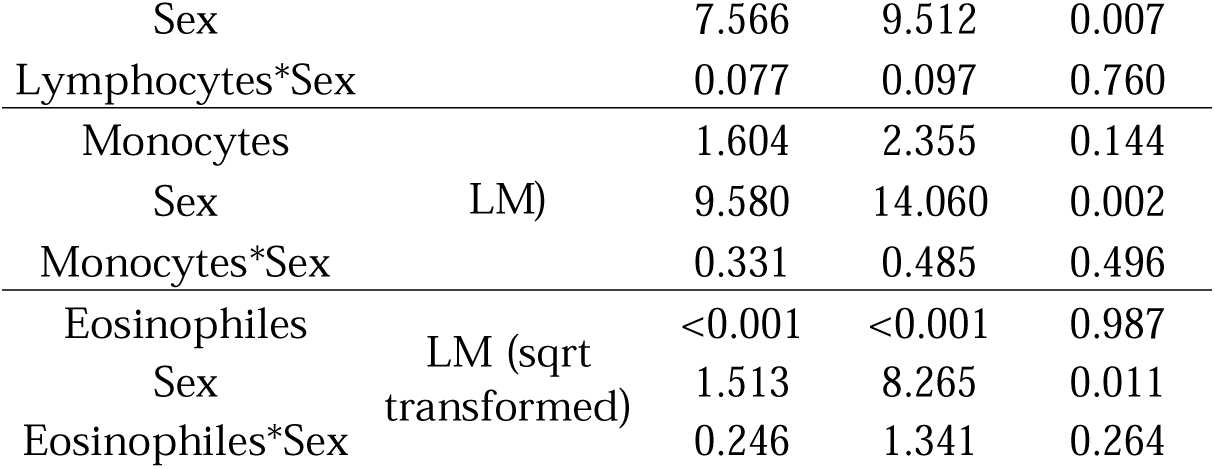

|  | Factors | Model | Sum Sq | F value | p-value |
| --- | --- | --- | --- | --- | --- |
| GFAP levels after 24 hours captivity | Pond salinity | Gamma GLM | 1.376 | 2.827 | 0.112 |
|  | Sex |  | 0.750 | 1.541 | 0.232 |
|  | Pond salinity*Sex |  | 0.552 | 1.135 | 0.303 |
|  | Osmolality | Gamma GLM | 0.117 | 0.225 | 0.642 |
|  | Sex |  | 1.720 | 3.301 | 0.088 |
|  | Osmolality*Sex |  | 0.669 | 1.284 | 0.274 |
|  | Hemoglobin-binding protein | Gamma GLM | 0.336 | 0.549 | 0.469 |
|  | Sex |  | 2.822 | 4.610 | 0.047 |
|  | Hemoglobin-binding protein*Sex |  | 0.074 | 0.121 | 0.733 |
|  | Neutrophiles | LM (log10 transformed) | <0.001 | <0.001 | 0.989 |
|  | Sex |  | 0.493 | 4.556 | 0.049 |
|  | Neutrophiles*Sex |  | 0.050 | 0.459 | 0.508 |
|  | Lymphocytes | LM (log10 transformed) | 0.156 | 1.608 | 0.223 |
|  | Sex |  | 0.682 | 7.024 | 0.017 |
|  | Lymphocytes*Sex |  | 0.072 | 0.744 | 0.401 |
|  | Monocytes | LM (log10 transformed) | 0.013 | 0.119 | 0.734 |
|  | Sex |  | 0.540 | 4.887 | 0.042 |
|  | Monocytes*Sex |  | 0.002 | 0.016 | 0.900 |
|  | Eosinophiles | LM | 1.859 | 2.795 | 0.114 |
|  | Sex |  | 3.829 | 5.756 | 0.029 |
|  | Eosinophiles*Sex |  | 0.025 | 0.037 | 0.849 |
| GFAP levels after 72 hours captivity | Pond salinity | LM | 0.491 | 0.657 | 0.430 |
|  | Sex |  | 8.536 | 11.430 | 0.004 |
|  | Pond salinity*Sex |  | 0.398 | 0.532 | 0.476 |
|  | Osmolality | LM | 0.420 | 0.550 | 0.469 |
|  | Sex |  | 8.246 | 10.790 | 0.005 |
|  | Osmolality*Sex |  | 0.189 | 0.247 | 0.626 |
|  | Hemoglobin-binding protein | Gamma GLM | 0.852 | 2.285 | 0.150 |
|  | Sex |  | 8.162 | 21.902 | <0.001 |
|  | Hemoglobin-binding protein*Sex |  | 0.035 | 0.094 | 0.763 |
|  | Neutrophiles | LM | 0.334 | 0.468 | 0.504 |
|  | Sex |  | 8.688 | 12.181 | 0.003 |
|  | Neutrophiles*Sex |  | 1.092 | 1.531 | 0.234 |
|  | Lymphocytes | LM | 0.035 | 0.043 | 0.838 |

|  |  |  |  |  |
| --- | --- | --- | --- | --- |
| Sex |  | 7.566 | 9.512 | 0.007 |
| Lymphocytes*Sex |  | 0.077 | 0.097 | 0.760 |
| Monocytes |  | 1.604 | 2.355 | 0.144 |
| Sex | LM) | 9.580 | 14.060 | 0.002 |
| Monocytes*Sex |  | 0.331 | 0.485 | 0.496 |
| Eosinophiles |  | <0.001 | <0.001 | 0.987 |
| Sex | LM (sqrt transformed) | 1.513 | 8.265 | 0.011 |
| Eosinophiles*Sex |  | 0.246 | 1.341 | 0.264 |

